# Non-cell autonomous inhibition of the Hedgehog response due to impaired cholesterol synthesis requires Ptch1/2 function

**DOI:** 10.1101/2020.12.10.420588

**Authors:** Carina Jägers, Henk Roelink

## Abstract

Congenital birth defects due to mutations of enzymes involved cholesterol synthesis, like Smith-Lemli-Opitz syndrome (SLOS) and Lathosterolosis are characterized by an accumulation of cholesterol precursors. The phenotype of both SLOS and Lathosterolosis have similarities to syndromes associated with abnormal Sonic hedgehog (Shh) signaling, consistent with the observation that cholesterol precursors and their derivative can inhibit Shh signaling. Two types of multipass membrane proteins play central roles in Shh signal transduction, the putative Resistance, Nodulation and Division (RND) antiporters Patched (Ptch)1 and −2, and the G-protein coupled receptor Smoothened (Smo). Sterols have been suggested as cargo for Ptch1/2, while Smo activity can affected both positively and negatively by steroidal molecules. We demonstrate that embryonic stem cells with mutations in the *7-dehydroxycholesterol reductase* (*7Dhcr*) or *sterol-C5-desaturase* (*Sc5d*) gene reduce the Hh response in adjacent wildtype cells when grown in mosaic organoids. This non-cell autonomous inhibitory activity of the mutant cells requires the presence of both Ptch1 and Ptch2. These observations support a model in which late cholesterol precursors that accumulate in cells lacking 7Dhcr are the cargo for Ptch1 and Ptch2 efflux activity and mediate the non-cell autonomous inhibition of Smo.

## Introduction

Hedgehog (Hh) signaling is critically important for embryonic development and patterning in most animals. Initial events in the transduction of the Hedgehog (Hh) signal include binding of the ligand to the 12-transmembrane hedgehog receptor Patched (Ptch) which releases a Ptch-mediated inhibition of the putative G-protein coupled receptor Smoothened (Smo) (Izzi et al., 2011; Marigo et al., 1996; Taipale et al., 2002). In the absence of Hh, Ptch inhibits Smo non-stoichiometrically (Taipale et al., 2002) and non-cell autonomously (Roberts et al., 2016) via a mechanism that remains poorly understood. Mammals have two Ptch homologs, Ptch1 and Ptch2 (Ptch1/2) that share overlapping functions (Alfaro et al., 2014; Lee et al., 2006; Roberts et al., 2016). Ptch1/2 are members of the Resistance-Nodulation Division (RND) family of antiporters that are present in all domains of life (Zhang et al., 2018). In bacteria, RND antiporters export their cargo in exchange for protons (Tseng et al., 1999) a function possibly conserved in mammals, although it is possible that Ptch1/2 can utilize other cation gradients (Myers et al., 2017). Bacterial RND transporters pump small lipophilic or amphiphilic molecules to the outside and eukaryotic RNDs plausibly have a similar function.

Although the precise nature of the cargo of Ptch1/2 antiporter activity remains unknown, several observations suggest that it is a steroidal Smoothened antagonist. The secretion of vitamin D3, a derivative of the cholesterol precursor 7-Dehydrocholesterol (7DHC), is Ptch1-dependent (Hausmann et al., 2009) and both vitamin D3 and 7DHC can repress Smo activity (Bijlsma et al., 2006; Linder et al., 2015). Several other known Smo modulators have a steroidal backbone, including the steroidal alkaloid cyclopamine (Incardona et al., 1998; Keeler, 1969; Sharpe et al., 2015). Sterols are derived from lanosterol, which is modified in a series of enzymatic reactions in the postsqualene pathway, culminating in the production of cholesterol or vitamin D (Bloch, 1965; Chojnacki and Dallner, 1988; Ernster and Dallner, 1995; Takeyama et al., 1997). Several congenital syndromes are caused by the loss of enzymes catalyzing the last steps in the cholesterol synthesis pathway (Figure 1A). Smith-Lemli-Opitz (SLO) is caused by mutations in the gene encoding 3β-hydroxysterol Δ7-reductase (7DHCR) (Fitzky et al., 1998), which catalyzes the reduction of the 7 double bond in 7-Dehydrodesmosterol (Kandutch and Russell, 1960). Phenotypically, SLOS is characterized by a distinctive facial appearance, cleft palate, microcephaly, a small, upturned nose, micrognathia, and ptosis. Furthermore, limb malformations like postaxial polydactyly, syndactyly of the second and third toe, and proximally placed thumbs are observed (SMITH et al., 1964). This syndrome has overlap with the malformations caused by altered Shh signaling. Biochemically, elevated levels of 7DHC and its isomer, 8-Dehydrocholesterol (8DHC) and decreased cholesterol are found in serum and tissue of SLO patients as well as in a mouse model (Tint et al., 1994), and increased 7DHC levels are used for diagnosis of SLO (Abuelo et al., 1995).

**Figure 1:**
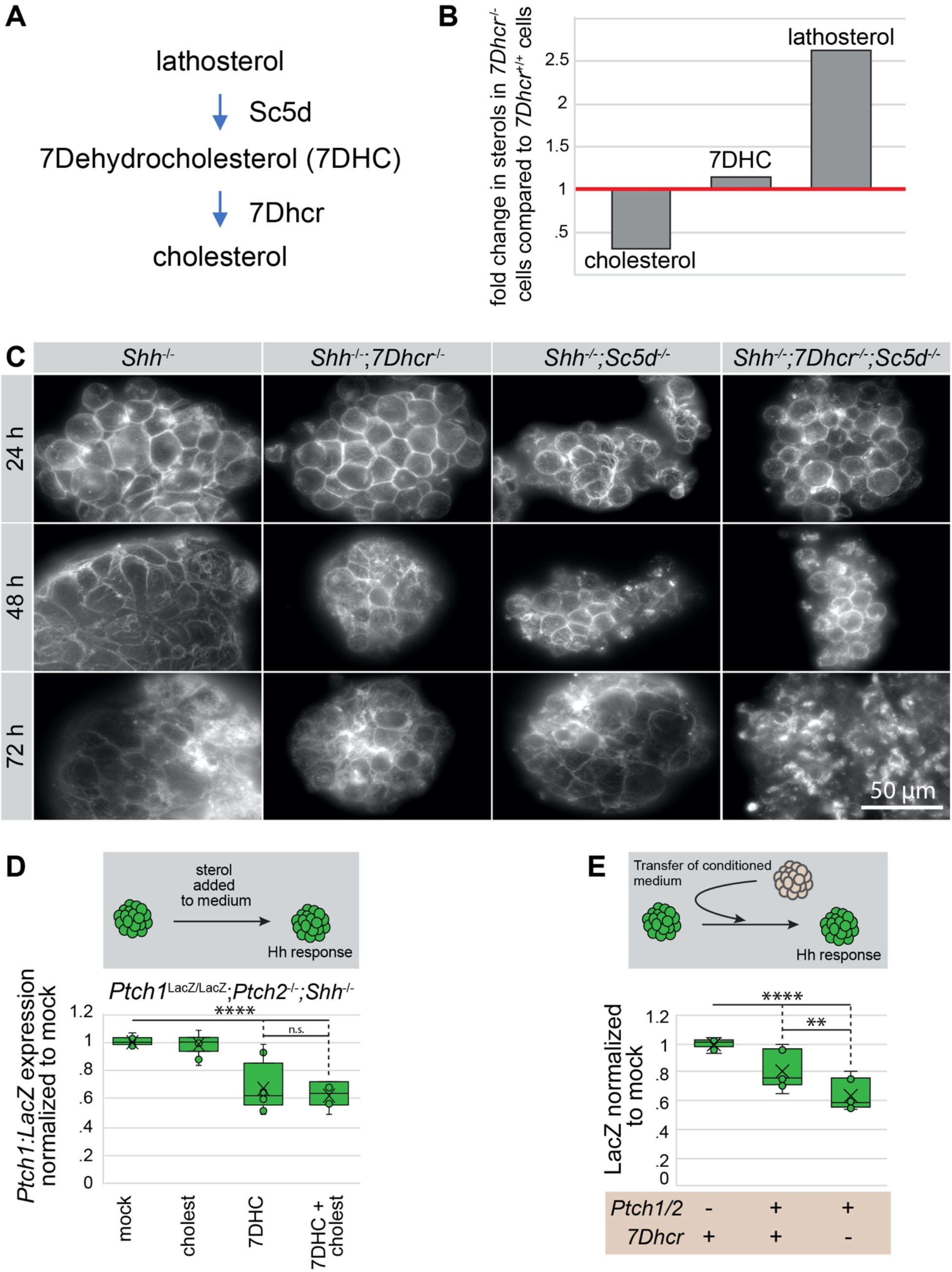
*Shh*^-/-^;*7Dhcr*^-/-^ inhibit the Hh response non-cell autonomously. **A:** Schematic of the last enzymatic reactions in cholesterol synthesis. Lathosterol is converted to 7DHC by the desaturase Sc5d, and 7DHC to cholesterol by the desaturase 7Dhcr. **B:** Cholesterol, 7DHC, and lathosterol levels were measured in *7Dhcr*^-/-^ cells 3 days after serum starvation by gas chromatography mass spectrometry (GC-MS) and compared to *7Dhcr^+/+^* cells. **C:** Staining of unesterified sterols in *Shh*^-/-^, *Shh*^-/-^;*7Dhcr*^-/-^, *Shh*^-/-^;*Sc5d*^-/-^, and *Shh*^-/-^;*7Dhcr*^-/-^;*Sc5d*^-/-^ SCOs with the fluorescent dye Filipin 3 days after serum starvation. **D:** *Ptch1:LacZ* measurement of *Ptch1*^LacZ/LacZ^;*Ptch2*^-/-^;*Shh*^-/^ reporter cells in SCOs that were treated with mock, 13 μM 7DHC, 13 μM cholesterol, or 13 μM 7DHC and 13 μM cholesterol for 3 days under serum starvation. Box-and-Whisker plot, n ≥ 3. **E:** *Ptch1:LacZ* measurement of *Ptch1*^LacZ/LacZ^;*Ptch2*^-/-^;*Shh*^-/^ reporter cells in SCOs that were cultured in medium conditioned by *Ptch1*^LacZ/LacZ^;*Ptch2*^-/-^;*Shh*^-/-^, *Shh*^-/-^, or *Shh*^-/-^;*7Dhcr*^-/-^ SCOs. Box-and-Whisker plot, n=3. \*\*\*\**p*<0.0001, \*\**p*<0.01, n.s. not significant.

Lathosterolosis is caused by mutations in the gene encoding the 3β-hydroxysteroid-Δ5-desaturase (SC5D) which desaturates lathosterol to 7DHC during cholesterol synthesis (Brunetti-Pierri et al., 2002). Symptoms of lathosterolosis have similarities with those observed in SLOS, including postaxial polydactyly, microcephaly, micrognathia, and toe syndactyly (Brunetti-Pierri et al., 2002). In a lathosterolosis mouse model, lathosterol levels are elevated 60 fold whereas levels of cholesterol were decreased similar to the SLOS mouse model (Krakowiak et al., 2003). Circulating maternal cholesterol can be transported to the fetus, making it unlikely that SLOS and lathosterolosis are predominantly caused by low fetal cholesterol, but instead it is plausible that the accumulation of late sterol precursors contributes to these birth defects, possibly by suppressing Hh signaling (Incardona et al., 1998).

Cell types in the developing neural tube arise in at stereotypic positions in response to Shh signaling. An effective in vitro model of the neural tube are spinal cord organoids (SCOs), which are differentiated aggregates of approximately 10^4^ embryonic stem cells (ESCs) (Wichterle et al., 2002). Mosaic SCOs can be generated by mixing cells with genotypically diverse cell lines in varying ratios. This approach allows us to address the non-cell autonomous effects caused by genetic alterations. Here, we provide evidence that loss of *7Dhcr* or *Sc5d* and associated accumulation of late sterol precursors can inhibit Shh signaling both cell-autonomously and noncell autonomously even in cells with an intact cholesterol synthesis pathway. The increased noncell autonomous inhibition requires the presence of Ptch1/2 function in those cells that accumulate the cholesterol precursors. These observations are consistent with a model that late sterol precursors are the cargo of Ptch1/2 antiporter activity in its ability to inhibit Smo non-cell autonomously.

## Results

### The loss of 7-Dehydrocholesterol reductase and/or Lathosterol 5-desaturase changes distribution of sterols

We have shown before that Ptch1/2 activity can inhibit Smo non-cell autonomously, supporting the notion that Ptch1/2 functions in the distribution and secretion of a Smo inhibitor (Roberts et al., 2016). To assess if this inhibitor could be a late sterol precursor, we made *Shh*^-/-^;*7Dhcr*^-/-^ cells and found that these cells have reduced amounts of cholesterol after three days of serum starvation as compared to the parental *Shh*^-/-^ cells (Figure 1B). To assess more global changes in the distribution sterols, we visualized the distribution of Filipin, a fluorescent stain of sterols (Blanchette-Mackie, 2000) used in the diagnosis of Niemann-Pick disease type C (Sokol et al., 1988). Similar to SLOS, the phenotype of lathosterolosis overlaps with that of congenital Shh malformations, raising the question if similar mechanisms impair Shh signal transduction. Besides *Shh*^-/-^;*7Dhcr*^-/-^ cells, we also stained *Shh*^-/-^;*Sc5d*^-/-^ cells as well as *Shh*^-/-^;*7Dhcr*^-/-^;*Sc5d*^-/-^ cells, cultured as SCOs in the absence of serum for up to 72 hours. *Shh^-/-^* cells had wrinkle-like accumulations of sterols in the plasma membrane on the first and second day of serum starvation (Figure 1C). However, few cells appeared with small rounder structures that accumulated on day 3. Cells lacking either *7Dhcr* or *Sc5d* had an initial Filipin staining pattern similar to *Shh*^-/-^ cells, while *Shh*^-/-^;*7Dhcr*^-/-^;*Sc5d*^-/-^ cells appeared smoother. At later timepoints, sterol distribution was different in *Shh*^-/-^;*7Dhcr*^-/-^;*Sc5d*^-/-^ cells with less sterol apparent on the plasma membrane and more pronounced staining in small domains, possibly endosomes. These results indicate that the *Sc5d* mutation is not strictly epistatic to the *7Dhcr* mutation, as might be predicted as Sc5d and 7Dhcr are thought to act in sequence in the cholesterol synthesis pathway. The finding that these mutations have an additive effect might suggest that alternate, parallel pathways function in cholesterol synthesis.

### Accumulation of late sterol precursors inhibits the Hh response

The loss of *7Dhcr* in SLOS is known to suppress Shh signaling, and a plausible hypothesis is that this is caused by the accumulation of a late sterol precursor that is inhibitory to Smo. To discern whether the accumulation of the cholesterol precursor 7DHC or the reduction of cholesterol can inhibit Shh signaling, SCOs constituting of *Ptch1*^LacZ/LacZ^;*Ptch2*^-/-^;*Shh*^-/-^ were treated with 13 μM 7DHC, 13 μM cholesterol, or a combination of both sterols. Activation of *Ptch1:LacZ*, measured by LacZ activity, was used as an indicator for the Hh pathway response (Figure 1D) (Goodrich et al., 1997). The reporter cells lack of both Ptch1 and Ptch2, resulting in an activation of Smo and elevated *Ptch1:LacZ* expression levels compared to heterozygous reporter cells (*Ptch1*^+/LacZ^;*Shh*^-/-^) (Alfaro et al., 2014; Roberts et al., 2016). Addition of 7DHC reduced the Shh pathway activity in *Ptch1*^LacZ/LacZ^;*Ptch2*^-/-^;*Shh*^-/-^ SCOs by 30 %. Cholesterol had no effect on the Hh response, neither when added alone nor in combination with 7DHC. This demonstrates that under these conditions 7DHC is an inhibitor of the Hh response, while cholesterol has no effect and cannot antagonize the presence of 7DHC. These results show that the inhibitory cholesterol precursor can be transferred via the culture medium. Therefore, we tested the inhibitory activity of conditioned supernatants. We transferred medium of SCOs consisting of wild type, *Ptch1/2*^-/-^ or *7Dhcr^-/-^* cells to *Ptch1*^LacZ/LacZ^;*Ptch2*^-/-^;*Shh*^-/-^ reporter cells and found that in addition to a Ptch-mediated inhibition, the medium conditioned by *7Dhcr*^-/-^ cells further decreased the Hh response in reporter cells by 40%, but not medium conditioned by cells lacking Ptch1/2 activity (Figure 1E). These results support the idea that a cholesterol precursor enriched in the medium via Ptch1/2 activity can inhibit the Hh response non-cell autonomously.

### Loss of 7-Dehydrocholesterol reductase and lathosterol 5-desaturase inhibit the Shh pathway non-cell autonomously in a Ptch1/2-dependent manner

To test if endogenous late cholesterol precursors like 7DHC or lathosterol can inhibit Shh signaling non-cell autonomously, we measured *Ptch1:LacZ* expression in *Ptch1*^LacZ/LacZ^;*Ptch2*^-/-^;*Shh*^-/^ reporter cells incorporated into mosaic SCOs that also contain 50% *Shh*^-/-^;*7Dhcr*^-/-^, *Shh*^-/-^;*Sc5d*^-/-^ or *Shh*^-/-^;*7Dhcr*^-/-^;*Sc5d*^-/-^ mESCs (Figure 2A). Both *Shh*^-/-^;*7Dhcr*^-/-^ and *Shh*^-/-^;*Sc5d*^-/-^ cells inhibited Shh pathway activity in the neighboring reporter cells (Figure 2B). The inhibition by *Shh*^-/-^;*7Dhcr*^-/-^ cells, however, was stronger than that by *Shh*^-/-^;*Sc5d*^-/-^ mESCs, suggesting that 7DHC is an important inhibitory sterol. In SCOs composed of equal amounts of *Ptch1*^LacZ/LacZ^;*Ptch2*^-/-^;*Shh*^-/-^ and *Shh*^-/-^;*7Dhcr*^-/-^;*Sc5d*^-/-^ mESCs the inhibition was stronger that that mediated by either *7Dhcr*^-/-^ or *Sc5d*^-/-^ cell alone. This additive effect is not easily reconciled with the idea of a linear pathway with 7Dhcr and Sc5d being enzymes acting in sequence but is consistent with and additive effect of these enzymes in sterol distribution (Figure 1C). The differences in the inhibitory potential of the different cell lines could not be explained by shifted ratios of reporter and repressor cells since the amount of CMFDA Green stained reporter cells remained the same in all 4 different types of mosaic nEBs 48 h after aggregation (Figure 2C). These results indicate that several early cholesterol precursors can inhibit Shh signaling non-cell autonomously. Moreover, the additive inhibitory effect of *Shh*^-/-^;*7Dhcr*^-/-^;*Sc5d*^-/-^ on the Shh pathway in not readily consistent the canonical linear cholesterol biosynthesis pathway.

**Figure 2:**
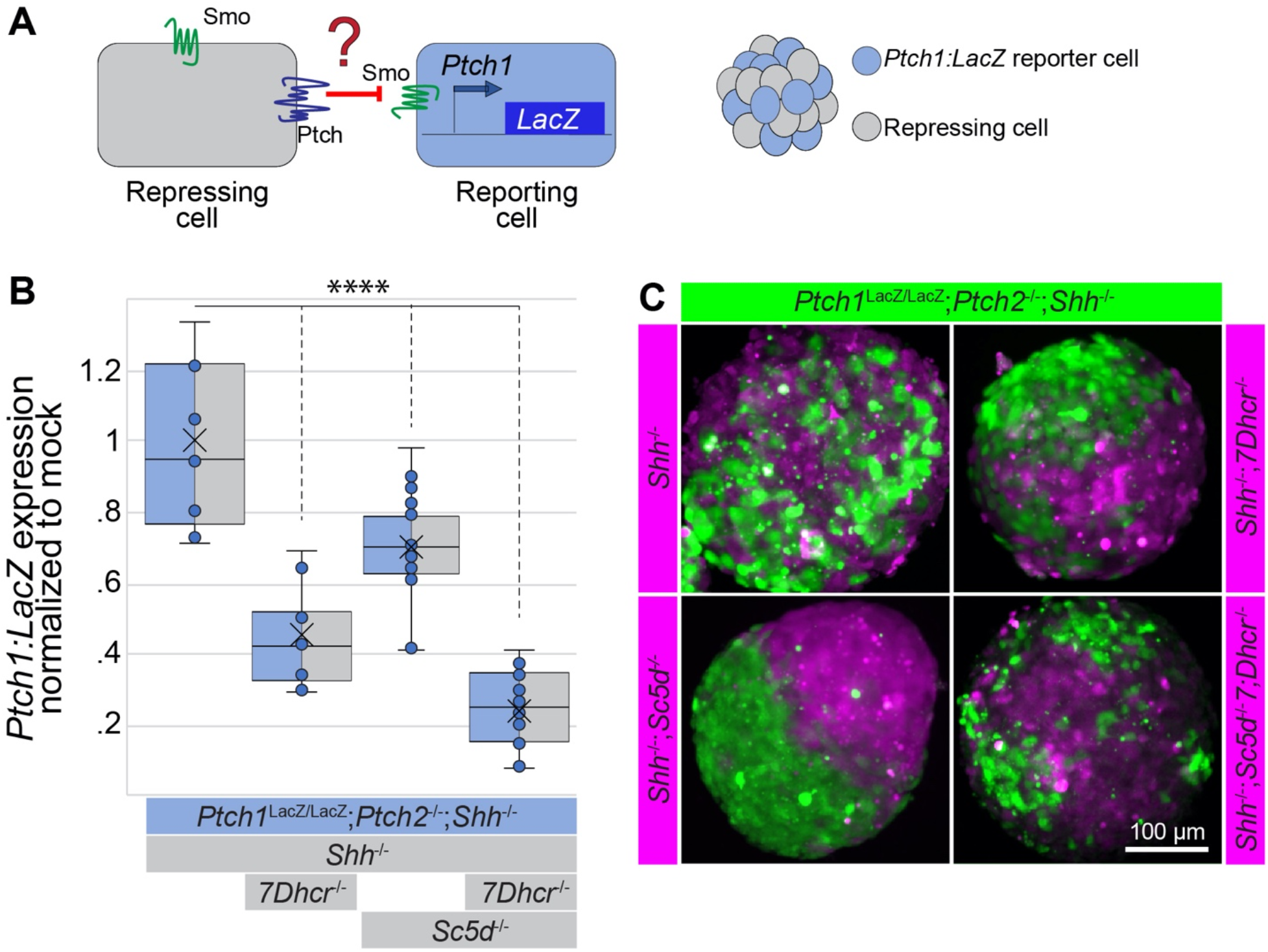
Non-cell autonomous inhibition of the Hh response by cells mutant for enzymes of cholesterol synthesis. **A:** Experimental setup: In mosaic SCOs consisting of equal amounts of reporter and repressor cells, repressor cells have Ptch1/2 but lack varying enzymes of cholesterol synthesis. The reporter cells with a *Ptch1:LacZ* reporter gene lack Ptch1/2 and therefore have an upregulated transcriptional Hh response. **B:** *Ptch1:LacZ* expression *Ptch1*^LacZ/LacZ^;*Ptch2*^-/-^;*Shh*^-/-^ reporter cells in mosaic SCOs after 3 days of serum starvation. Box-and-Whisker plot, n=5, **** *p*<0.0001. **C:** Fluorescent images of mosaic SCOs consisting of reporter and repressor cells (genotypes indicated) that were loaded with lineage tracing dyes (CellTracker^™^ Green and Blue). Images are pseudo-colored.

### Ptch1/2 activity is required for non-cell autonomous inhibition

A central question that remains is if Ptch1/2 function is required of the non-cell autonomous inhibition of the Hh response by late sterol precursors. We compared the response to 7DHC in SCOs consisting of 100% *Ptch1*^LacZ/LacZ^;*Ptch2*^-/-^;*Shh*^-/-^ cells to mosaic nEBs consisting of 50% *Ptch1*^LacZ/LacZ^;*Ptch2*^-/-^;*Shh*^-/-^ reporter and 50% Ptch1/2-containing cells (Figure 3A). Exogenously applied 7DHC decreased the Hh response in *Ptch1*^LacZ/LacZ^;*Ptch2*^-/-^;*Shh*^-/-^ reporter cells to a much greater extend when Ptch1/2-containing cells were present in the same SCO, compounding the inhibitory function of Ptch1/2 alone. This suggests that 7DHC from the medium can be acted upon, and potentiated by Ptch1/2 function to inhibit the Hh response non-cell autonomously. Ptch1/2 function was unable to potentiate Smo antagonist cyclopamine, suggesting that cyclopamine inhibits Smo directly independent of Ptch1/2 function (Figure S1).

**Figure 3:**
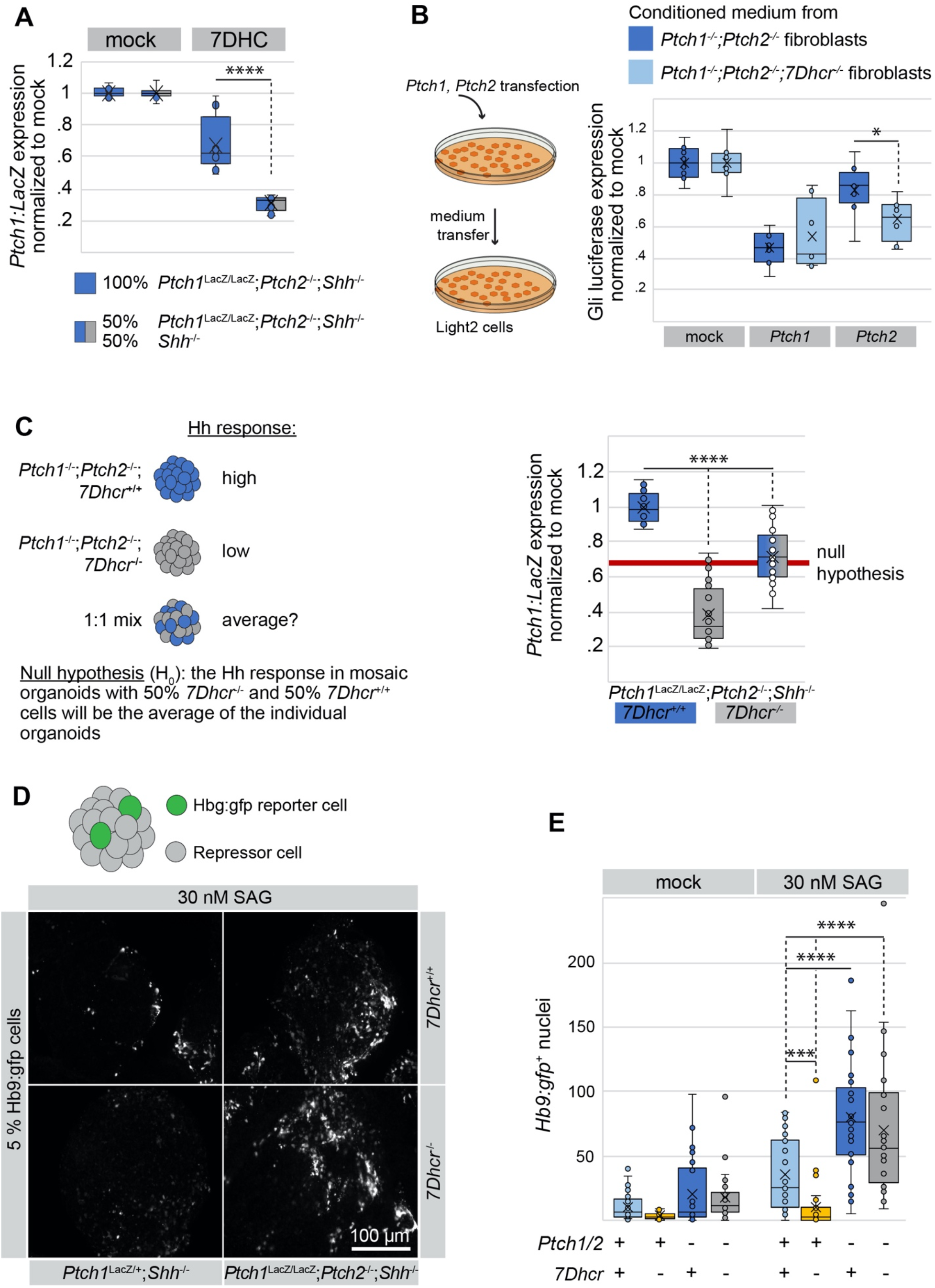
Non-cell autonomous inhibition by late cholesterol precursors requires Ptch1/2. **A:** *Ptch1:LacZ* measurement of *Ptch1*^LacZ/LacZ^;*Ptch2*^-/-^;*Shh*^-/-^ reporter cells in SCOs or in mosaic SCOs with 50 % *Shh*^-/-^ mESCs after 3 days of serum starvation. 13 μM 7DHC was added after the first night of SCO aggregation. Ptch1:LacZ levels are normalized to the respective mock treatment to account for lower numbers of reporter cells and inhibition by Ptch1/2 in mosaic SCOs. Box-and-Whisker plot, n=4. **B:** Experimental setup and Gli luciferase measurements of Hh responsive Light2 cells. *Ptch1*^-/-^;*Ptch2*^-/-^ or *Ptch1*^-/-^;*Ptch2*^-/-^;*7Dhcr*^-/-^ fibroblasts were transfected with mock, *Ptch1*, or *Ptch2* and their conditioned medium was filtered and transferred onto Light2 cells with 30 nM SAG. Gli luciferase levels were normalized to Renilla luciferase and all conditions to their respective mock transfection. Box-and-Whisker plot, n=4. **C:** Graphic representation of the null hypothesis (indicated by red line): If Ptch1/2 is not required for non-cell autonomous inhibition, *Ptch1:LacZ* expression in mosaic SCOs will be the average of the individual SCOs. *Ptch1:LacZ* measurement of *Ptch1*^LacZ/LacZ^;*Ptch2*^-/-^;*Shh*^-/-^, *Ptch1*^LacZ/LacZ^;*Ptch2*^-/-^;*Shh*^-/-^*7Dhcr*^-/-^, and mosaic SCO consisting of equal amounts of the two reporter lines 3 days after serum starvation. Box-and-Whisker plot, n=4. **D:** Mosaic SCOs with 5% *Hb9:gfp* reporter cells and the indicated repressor cells were serum starved for 3 days and treated with 30 nM SAG for the last 24 hours. SCOs were fixed and *Hb9:gfp* expression imaged. Shown are representative images of mosaic SCOs for each condition. **E:** Quantification of *Hb9:gfp* positive nuclei of the experiment shown in D, shown in a Box-and-Whisker plot, n=3.* *p*<0.05, *** *p*<0.001, ****p<0.0001

We tested if Ptch1/2 is sufficient for non-cell autonomous inhibition by cells mutant for *7Dhcr* or *Sc5d*. We expressed Ptch1 or Ptch2 in *Ptch*1/2- or *7Dhcr*-deficient fibroblasts (*Ptch1*^LacZ/LacZ^;*Ptch2*^-/-^ and *Ptch1*^LacZ/LacZ^; *Ptch2*^-/-^; *7Dhcr*^-/-^) and transferred their supernatant onto Light2 reporter cells. Conditioned medium from Ptch1- and Ptch2-expressing cells inhibited the Shh pathway response in the presence of SAG equally and independent of 7Dhcr in the conditioning cells (Figure 3B).

To further assess the role of Ptch1/2 in non-cell autonomous inhibition of Shh signaling by cholesterol precursors, we mixed Ptch1/2-deficient *Ptch1*^LacZ/LacZ^;*Ptch2*^-/-^;*Shh*^-/-^ and *Ptch1*^LacZ/LacZ^;*Ptch2*^-/-^;*Shh*^-/-^;*Dhcr7*^-/-^ SCOs and compared their *Ptch1:LacZ* expression to nEBs consisting of either genotype alone. As expected, the Hh response is constitutively high in *Ptch1*^LacZ/LacZ^;*Ptch2*^-/-^;*Shh*^-/-^ cells. However, the loss of *7Dhcr* decreases Ptch1:LacZ expression by approximately 70% (Figure 3C). If Ptch1/2 are indeed required for non-cell autonomous Shh pathway inhibition, 7Dhcr- and Ptch1/2 deficient cells should not be able to inhibit the Shh pathway response and the *Ptch1:LacZ* expression of the mosaic nEBs should be exactly half of the difference between *Ptch1*^LacZ/LacZ^;*Ptch2*^-/-^;*Shh*^-/-^ and *Ptch1*^LacZ/LacZ^;*Ptch2*^-/-^;*Shh*^-/-^;*7Dhcr*^-/-^ nEBs. The null hypothesis would therefore be H_0_ = 1 – ½ (average *Ptch1*^LacZ/LacZ^;*Ptch2*^-/-^;*Shh*^-/-^- average *Ptch1*^LacZ/LacZ^;*Ptch2*^-/-^;*Shh*^-/-^;*7Dhcr*^-/-^), indicated by a red line in Figure 3C. We found that the overall Hh response in *Ptch1/2*-deficient mosaic SCOs was nearly the same as H_0_, supporting the hypothesis that cells lacking Ptch1/2 function cannot inhibit the Hh response non-cell autonomously.

To further discern the role of Ptch1/2 in repressing and responding cells, we mixed 5% *Hb9:gfp* mESCs (Wichterle et al., 2002) with a panel of *Ptch1/2*-proficient/deficient and/or *7Dhcr*-proficient/deficient mESCs. *Hb9:gfp* mESCs carry a transgene in which the Hh-sensitive *Hb9* promoter drives *gfp*, and thus serve as reporters for activation of the Hh response (Figure 3D). The Hh pathway response was activated in the reporter cells with the Smo agonist SAG and the inhibitory potential of the interspersed cells was assessed by counting the number of *Hb9:gfp*-positive nuclei as a measure of Hh pathway activation in SCOs (Wichterle et al., 2002). Compared to uninhibited *Hb9:gfp* reporter cells in Ptch1/2-deficient mosaic nEBs, Ptch1/2 alone decreased the number of positive nuclei by more than half. Consistent with our results described earlier, Ptch1/2 containing but 7Dhcr deficient cells decreased the number of positive nuclei even further. In the absence of Ptch1/2 and 7Dhcr, however, Hb9:gfp expression was uninhibited, similar to the level of Ptch1/2-deficient mosaic nEBs. This demonstrates that a loss of Ptch1/2 activity is epistatic to a loss of 7Dhcr function in repressing cells and is consistent with the notion that Ptch1/2 is required for non-cell autonomous inhibition. As the addition of SAG increased the number of *Hb9:gfp* positive nuclei in every condition, the inhibitory factor in the *7Dhcr*^-/-^ cells is possibly a competitive inhibitor to SAG, further indicating that Smo is the target of 7DHC inhibition of the Hh response (Figure 3E).

## Discussion

In humans, mutations in the *DHCR7* gene that codes for 3β-hydroxysteroid *Δ*^7^-reductase are associated with the Smith-Lemli-Opitz Syndrome (SLOS), a congenital malformation syndrome (Fitzky et al., 1998). Some of the SLOS-associated malformations resemble birth defects observed in animals with reduced Hh signaling (Roessler et al., 1996), raising the hypothesis that the accumulating cholesterol precursor 7DHC reduces Shh signaling. Here, *Shh*^-/-^;*7Dhcr*^-/-^ mESCs were found to inhibit the Shh pathway non-cell autonomously. Importantly, Ptch1/2 was found to enhance the inhibitory effect of exogenously added 7DHC on reporter cells devoid of Ptch1/2, arguing for 7DHC being an endogenous cargo of Ptch1/2. Furthermore, Ptch1/2 are required for non-cell autonomous inhibition as *7Dchr*^-/-^ cells also deficient for Ptch1/2 were not able to inhibit the Hh response in adjacent cells.

### Possible scenario for the Ptch1/2-dependent inhibition of Smo by cholesterol precursors

We propose the following model based on the data presented here (Figure 4): Ptch1/2 is able to inhibit Smo non-cell autonomously and its inhibitory function is even enhanced when the cells are deficient for one or more enzymes of the cholesterol synthesis pathway. This is likely due to the Ptch1/2-mediated transport of inhibitory cholesterol precursors like 7DHC, increasing their inhibitory potential by a yet unknown mechanism (Bijlsma et al., 2006; Linder et al., 2015). To this point, we cannot exclude that other sterols that accumulate in the absence of 7Dhcr or Sc5d are the preferred cargo of Ptch1/2 that inhibits Smo.

**Figure 4:**
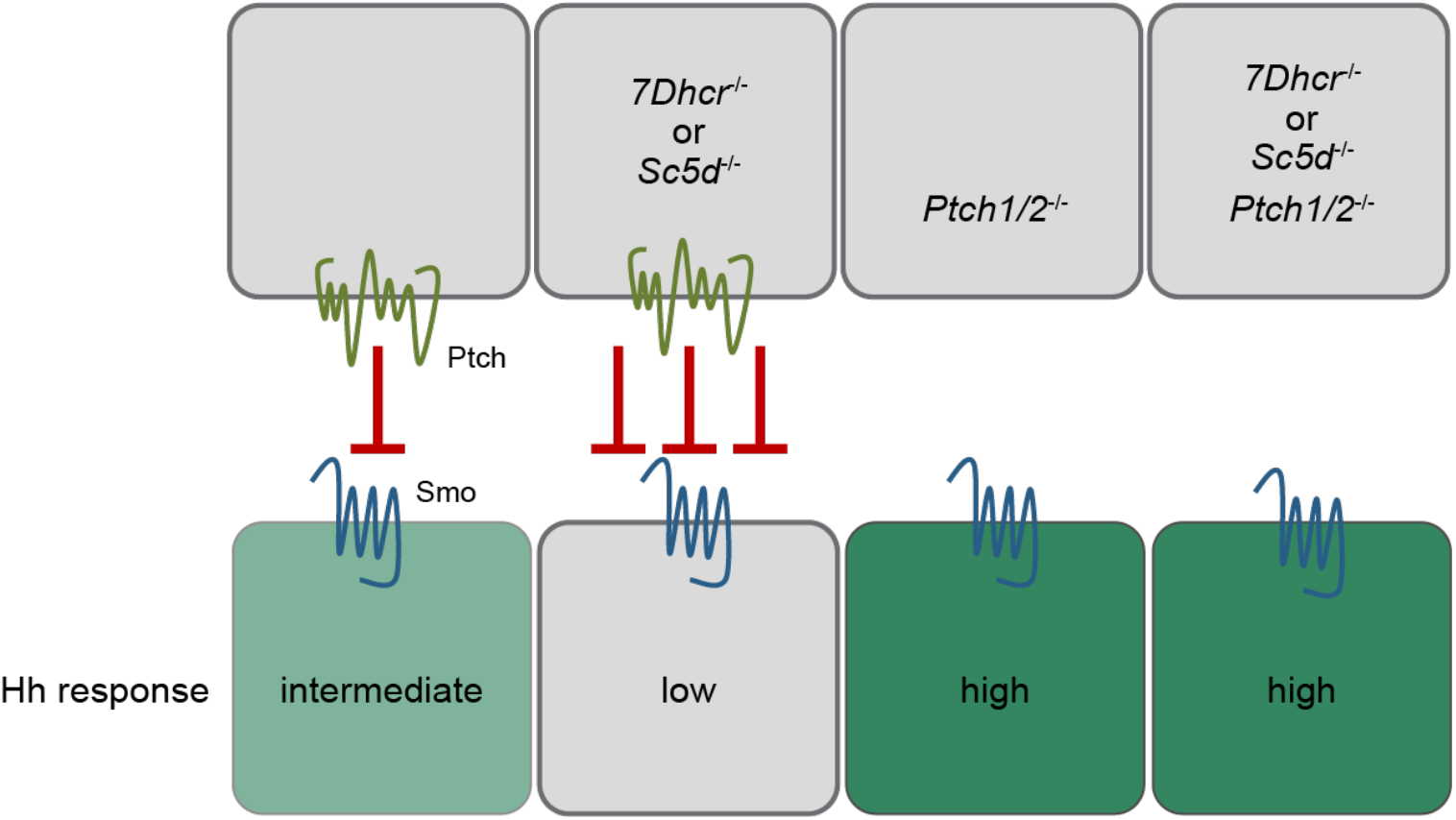
Model of non-cell autonomous Smo inhibition by Ptch1/2. Ptch1/2 inhibits Smo non-cell autonomously and does so more potently when the Ptch1/2-containing cells lack enzymes of cholesterol synthesis, resulting in a decreased transcriptional Hh response. In the absence of Ptch1/2 and independent of the absence or presence of enzymes of cholesterol synthesis, Smo in uninhibited and the transcriptional Hh response is high.

### Does Ptch transport a Smo-inhibitory sterol?

There are many Smo-modulating small molecules, both endogenous and exogenous. Several of the molecules affecting Smo activity have a steroidal structure. Together with evidence that Ptch1/2 can inhibit Smo non-cell autonomously, suggests that in the absence of the Hh ligand, Ptch transports a sterol that inhibits Smo. Cyclopamine, a steroidal alkaloid inhibits Smo via direct binding (Chen et al., 2002). However, we find that Ptch1/2 does not exacerbate Smo inhibition by cyclopamine (Figure S1), arguing against that this steroidal alkaloid is a cargo of Ptch1/2. The ubiquity of cholesterol makes it an unlikely substrate for the Ptch1/2 function in Smo inhibition. However, late cholesterol precursors are good candidates. Vitamin D3 is derived from 7DHC via photolysis (Holick and Clark, 1978). When 7DHC accumulates in the absence of 7Dhcr, the synthesis of vitamin D3 is increased (Prabhu et al., 2016). Vitamin D3/cholecalciferol, the precursor of vitamin D3, is capable of inhibiting Smo more efficiently than 7DHC and is secreted by Ptch1 (Bijlsma et al., 2006). Furthermore, vitamin D3, was reported to be released into the extracellular space by Ptch1 and efficiently inhibit the translocation of Smo to the primary cilium (Linder et al., 2015).

### Evidence for alternative pathways in the cholesterol biosynthesis pathway

Cholesterol synthesis begins with acetyl-CoA and has a multiplicity of intermediates, each requiring catalyzation by unique enzymes to progress to the next precursor molecule. Kandutsch and Russell proposed a linear but bifurcated pathway in 1960 for the synthesis of cholesterol that only differ in an additional double bond in the alkyl side chain appended to the CD-ring. This double bond can be resolved at any time by the 3β-hydroxysterol Δ^24^-reductase. Sc5d acts upstream of 7Dhcr and a mutation would therefore be predicted to be epistatic to a mutation in the *7Dhcr* gene. Contrary, we observed an additive inhibitory effect of the two mutations in combination on the Shh pathway. Thus, additional functions of these enzymes or additional, parallel pathways in the final steps of cholesterol synthesis are conceivable. Porter et. al proposed, with reservation, that 8-Dehydrocholesterol (8DHC), an isomer of 7DHC, can be synthesized from the precursor of lathosterol (cholesta-8(9)-en-3β-ol) by Sc5d (Porter, 2002; Porter and Herman, 2011). 8DHC and 7DHC differ in the position of a double bond that can be swapped in both directions by the enzyme that synthesizes lathosterol from its precursor, 3β-hydroxysterol Δ8,Δ7-isomerase (EBP). Although this alternative pathway does not explain the additive inhibitory effect of combinatorial *Sc5d* and *7Dhcr* knock-outs - a lack of Sc5d would still terminate the synthesis of cholesterol at the level of lathosterol - it might suggest that the cholesterol synthesis pathway is not strictly linear and that enzymes of cholesterol synthesis can exhibit additional functions.

### The mechanism of Smo inhibition by Ptch1/2

The hypothesis that the non-cell autonomous inhibition of the Shh pathway by Ptch1/2 is mediated by its antiporter function originates from the observation that a mutation in the antiporter channel (Ptch1D499A) blocks Ptch1 activity and subsequently, the Hh response (Alfaro et al., 2014; Taipale et al., 2002). Further evidence of this model came from the observation that the Ptch1/2-mediated inhibition of Smo is non-cell autonomous (Roberts et al., 2016).

As a member of the RND family of proton driven antiporters Ptch1/2 is thought to require a proton gradient to transport molecules across membranes (Tseng et al., 1999). Sodium has been proposed as an alternative ion that would be transported across the cell membrane in exchange for the Smo inhibitory molecule. Besides the question of “what” Ptch1/2 transports, the “how/where” has been the point of more recent discussion. The Ptch1 monomer revealed a hydrophobic pore-like structure in cryo-electron microscopy (cryo-EM) models that can be occupied by the N-terminus of Shh. In bacterial RNDs this is the channel that lets in protons and blocking this channel by Shh would provide a simple mechanism by which Shh blocks Ptch1/2 function. Dispatched, another member of the RND superfamily, is a homotrimer (Etheridge et al., 2010), as is *Drosophila* Ptch (Lu et al., 2006) indicating that also Ptch1 functions as a trimer. In bacterial RNDs trimerization forms a central channel through which secreted cargo leaves the cell, and that Ptch would function like bacterial RNDs, something indicated by conservation in their structures.

## Materials and Methods

### CRISPR/Cas9 mediated mutagenesis

The CRISPR/Cas9 genome editing technique was used to knock out the *7Dhcr* and *Sc5d* gene (Ran et al. 2013). The pX458/ pSpCas9(BB)-2A-Puro and pX459/ pSpCas9(BB)-2A-GFP expression plasmids were a gift from the Doudna lab (University of California, Berkeley, USA). sgRNAs were designed using the online tool provided by the Zhang lab (http://tools.genome-engineering.org, MIT, Massachusetts, USA) and ordered from IDT DNA Technologies (Iowa, USA). Cloning of sgRNA and Cas9 expressing plasmids was performed according to (Ran et al., 2013)

### Tissue culture

*Ptch1*^LacZ/LacZ^;*Ptch2*^-/-^;*Shh*^-/-^ and *Shh*^-/-^ mESCs were generated using TALEN mediated mutagenesis and are described elsewhere (Roberts et al., 2016). The *Ptch1*^LacZ/LacZ^;*Ptch2*^-/-^;*Shh*^-/-^ line originates from the *Ptch1*^LacZ/LacZ^ line (Goodrich et al., 1997). All cell lines were cultured under standard conditions in ES medium (DMEM/Gibco, 15 % FBS/Gibco, 2mM L-glutamine/Thermo Fisher Scientific, 1X Penicillin Streptomycin/Thermo Fisher Scientific, 1X MEM Non-Essential Amino Acids Solution/Thermo Fisher Scientific, 1X Nucleosides for ES cells/EMD Millipore, 1X ß-Mercaptoethanol, 1X GST-LIF/Millipore) in tissue culture dishes coated with gelatin.

For generation of SCOs, 2 x 10^5^ – 4 x 10^5^ mESCs were cultured in 60mm Petri dishes in DFNB medium (Neurobasal medium/Gibco, DMEM F12 1:1/Gibco, 1.5 mM L-glutamine/Thermo Fisher Scientific, 1X Penicillin Streptomycin/Thermo Fisher Scientific, 1X B27^®^ Supplement/Gibco, 0.1 mM ß-Mercaptoethanol) on a rotating platform. After 48 h, 10 μM Retinoic Acid (Sigma/Aldrich) was added.

7DHC, cholesterol, and lathosterol (all Sigma/Aldrich) were dissolved in ethanol and used at 13 μM. All reagents were added directly to aggregating SCOs.

Fibroblasts were grown in DMEM + 10% FBS (Atlas Biologicals) and transfected with Lipofectamine2000 according to the manufacturer’s instructions. The medium was exchanged with DMEM 24h after transfection for medium conditioning (24h). The conditioned medium was then removed, filtered, and transferred onto Light2 cells for another 48h with 30 nM SAG. Gli- and Renilla-luciferase levels in the Light2 cells were measured with the Dual luciferase kit according to the manufacturer’s instructions. Gli luciferase/Renilla luciferase levels were normalized to the respective mock-transfected control.

### Cell tracking

For cell lineage tracking experiments, cells were incubated with 500 μM CMFDA Green or CMAC Blue (both Invitrogen) in DFNB medium for 45 min at 37 °C and seeded as described above. After 48 h (or respective time points), SCOs were fixed in 4 % paraformaldehyde (Thermo Fisher Scientific) in 1X PBS for 10 min and mounted directly in Fluoromount G Mounting Medium (Thermo Fisher Scientific). Images were taken with a Zeiss Observer fluorescence microscope with a 20X and 10X objective.

### Filipin staining

For filipin staining, SCOs were fixed in 4 % paraformaldehyde (Thermo Fisher Scientific) in 1X PBS for 10 min after one, two, or three days. SCOs were stained in 30 μM filipin III (Sigma/Aldrich) in 1X PBS for 30 min in the dark and mounted in Fluoromount G Mounting Medium (Thermo Fisher Scientific) after washing once with 1X PBS. Images were taken with a Zeiss Observer fluorescence microscope with a 63X objective and oil immersion.

### Reporter Gene Assay for Ptch1:LacZ induction

*Ptch1:LacZ* expression levels were quantified 72 h after aggregation of SCOS using the Galacto-Light Plus^™^ system (Applied Biosciences) according to the manufacturer’s instructions. Shortly, SCOs were collected in plastic tubes, washed with 1X PBS, and lysed using lysis buffer (100 mM potassium phosphate pH 7.8, 0.2% Triton X-100). Lysates were incubated with 70 μl Reaction buffer for 30 min in a 96 well plate, followed by an incubation with Accelerator-II for 15 min. Signal was read in a microplate luminometer for 5 s per well. *Ptch1:LacZ* levels were normalized to total protein using the Bradford reagent (BioRad).

### Statistics

Single Factor ANOVA was used to analyze more than two conditions, followed by a Student’s *t*-test with a two-tailed distribution assuming unequal variance comparing two conditions. \**p*<0.05, \*\**p*<0.01, \*\*\**p*<0.001, \*\*\*\**p*<0.0001

## Acknowledgements

We thank Dr. Anthony T. Lavarone at the QB3/Chemistry Mass Spectrometry Facility (UC Berkeley) for help with obtaining the mass spectrometry data.

## Author contributions

All experiments were performed by CJ. Experiments were designed by CJ and HR. The manuscript was written by CJ and HR.

---

**Figure S1:**
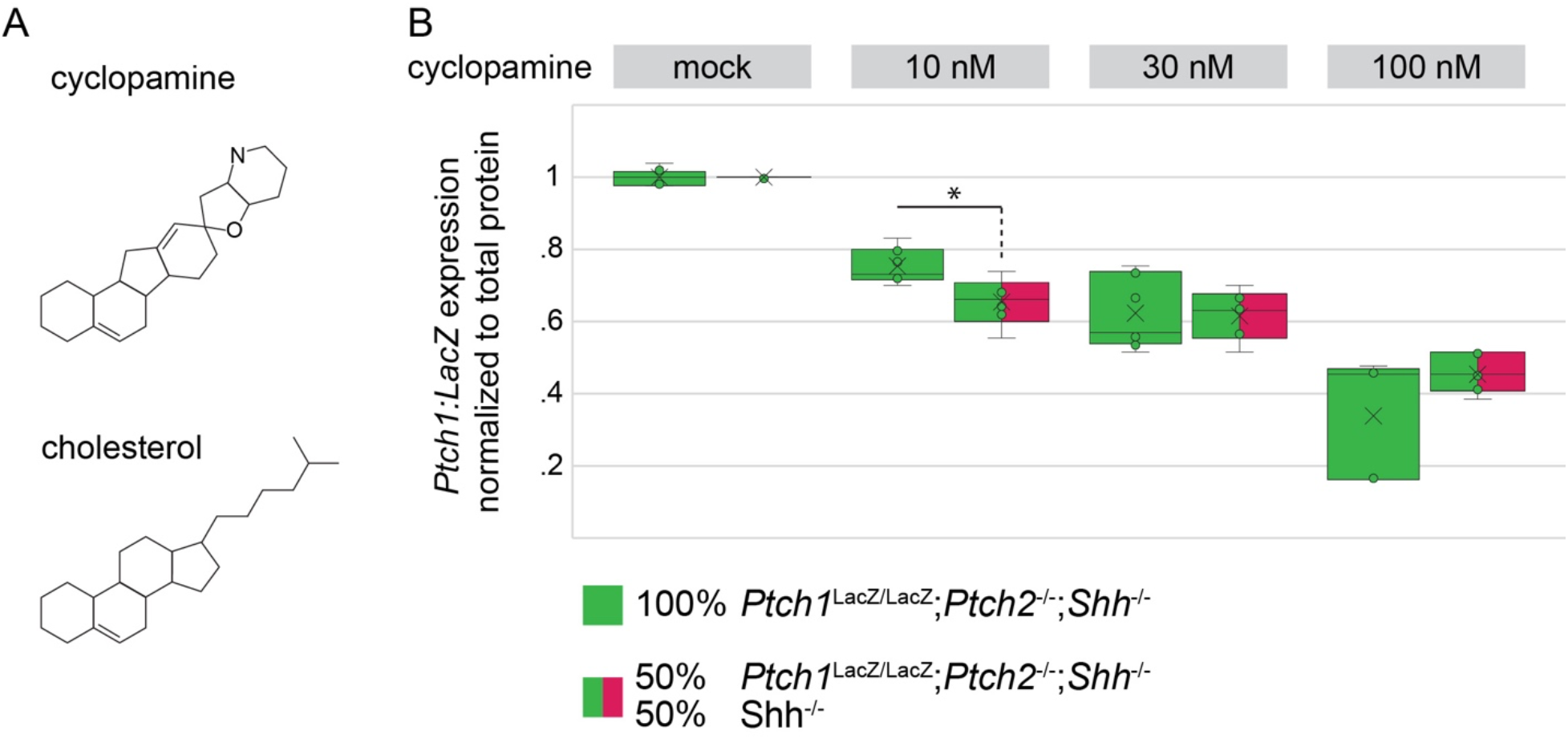
Ptch1/2 does not enhance the inhibition of Smo by cyclopamine. **A:** Structural comparison of the steroidal molecules cyclopamine and cholesterol. **B:** *Ptch1:LacZ* measurement of *Ptch1*^LacZ/LacZ^;*Ptch2*^-/-^;*Shh*^-/-^ reporter cells in SCOs or in mosaic SCOs with 50 % *Shh*^-/-^ mESCs after 3 days of serum starvation. The indicated concentrations of cyclopamine were added after the first night of SCO aggregation. *Ptch1:LacZ* levels are normalized to the respective mock treatment to account for lower numbers of reporter cells and inhibition by Ptch1/2 in mosaic SCOs. Box-and-Whisker plot, n=6, \**p*<0.05.

